# Simultaneous B and T cell acute lymphoblastic leukemias in zebrafish driven by transgenic MYC: implications for oncogenesis and lymphopoiesis

**DOI:** 10.1101/281469

**Authors:** Chiara Borga, Gilseung Park, Clay Foster, Jessica Burroughs-Garcia, Matteo Marchesin, Syed T. Ahmed, Silvia Bresolin, Lance Batchelor, Teresa Scordino, Rodney R. Miles, Geertruy te Kronnie, James L. Regens, J. Kimble Frazer

## Abstract

Precursor-B cell acute lymphoblastic leukemia (pre-B ALL) is the most common pediatric cancer, but there are no useful zebrafish pre-B ALL models. We describe the first highly-penetrant zebrafish pre-B ALL, driven by human *MYC*. Leukemias express B lymphoblast-specific genes and are distinct from T cell ALL (T-ALL)—which these fish also develop. Zebrafish pre-B ALL shares *in vivo* features and expression profiles with human pre-B ALL, and these profiles differ from zebrafish T-ALL or normal B and T cells. These animals also exhibit aberrant lymphocyte development. As the only robust zebrafish pre-B ALL model and only example where T-ALL also develops, this model can reveal differences between *MYC*-driven pre-B vs. T-ALL and be exploited to discover novel pre-B ALL therapies.

**Statement of significance:** We describe the first robust zebrafish pre-B ALL model in MYC-transgenic animals known to develop T-ALL, revealing the only animal model with both human ALL types. We also describe aberrant multi-lineage lymphopoiesis. This powerful system can be used to study *MYC*-driven leukemogenesis and discover new pre-B ALL targeted therapies.

## Introduction

Acute lymphocytic leukemia (ALL), a common cancer and the most prevalent childhood malignancy, comprises >25% of pediatric neoplasia in the U.S., with ∼85% being pre-B ALL (1, 2). Relapses are all-too-common, making ALL the highest cause of pediatric cancer-related death (3). Thus, there is a dire need for animal models of pre-B ALL to identify new molecular targets and discover new therapies, but efforts are impeded by a lack of *in vivo* models amenable to genetic and drug screens.

Zebrafish (*Danio rerio*) provide one potential solution, since they can model human leukemias accurately (4), have practical advantages (genetic tractability, high-throughput screens, cost), and share hematopoietic, oncogenic, and tumor suppressive pathways with humans (5). These features permitted the creation of several zebrafish T cell ALL (T-ALL) models that mimic the human disease (6-10), which subsequently led to key findings in T-ALL genetics, disease progression mechanisms, and signaling (11-16), as well as facilitating screens for new treatment agents (17, 18). However, despite the even greater clinical impact of pre-B ALL, effective zebrafish models lag behind. A single report of zebrafish B-ALL using transgenic *ETV6-RUNX1* (19) had low penetrance and long latency (∼3% by 1 year), and no subsequent reports with this or any other B-ALL model exist.

Here we utilized a cell-specific fluorophore, *lck*:*eGFP (20),* that labels zebrafish B and T cells differentially to discover the first robust *D. rerio* B-ALL model. Surprisingly, B-ALLs occur in an already-established T-ALL model driven by transgenic human *MYC* (10), and they are so prevalent that many animals actually have coincident B-and T-ALL. An intensive investigation of this new model using several approaches revealed a number of important findings. First, *hMYC*-induced B-ALL are pre-B subtype, express immature B cell transcripts, and like human pre-B ALL, spread aggressively to lymphoid and non-lymphoid tissues. Second, pre-B ALL express low levels of *lck*, and thus are dimly-fluorescent in these animals, unlike the brightly-fluorescent T-ALL of this model. Low *LCK* expression is conserved in human pre-B ALL. Third, in addition to their differential *lck*:*eGFP* expression, we report a two-gene classifier that distinguishes pre-B from T-ALL in *hMYC* fish. Finally, expression profiles of zebrafish pre-B ALL, T-ALL, and normal B and T cells revealed abnormal lymphopoiesis that may underlie the molecular pathogenesis of *hMYC*-driven ALL. In summary, we report a novel and robust pre-B ALL model, the first in zebrafish. Besides its value for genetic and drug screens, to our knowledge, *hMYC* fish represent the only animal model that develops both pre-B and T-ALL, providing a unique tool to explore molecular mechanisms of both human ALL types in the same genetic context, or even the same animal.

## Results

### Human *MYC* induces two zebrafish ALL types with distinct expression signatures

Mammalian *Myc*/*MYC* transgenes driven by a *D. rerio rag2* promoter induce zebrafish T-ALL (6, 10). To detect and monitor ALL progression, we built double-transgenic fish by crossing Tg(*rag2:hMYC*) to Tg(*lck:eGFP*) fish, where a zebrafish *lck* promoter controls GFP expression (20). Henceforth, we refer to this double-transgenic line as *hMYC;GFP*. To study T-ALL in this system, we performed RNA microarray on FACS-purified GFP^+^ ALL cells, analyzing 10 *hMYC;GFP* malignancies and 3 cancers from *hlk* fish (9), another zebrafish T-ALL model (see Fig. 1A for example animals).

**Fig. 1.**
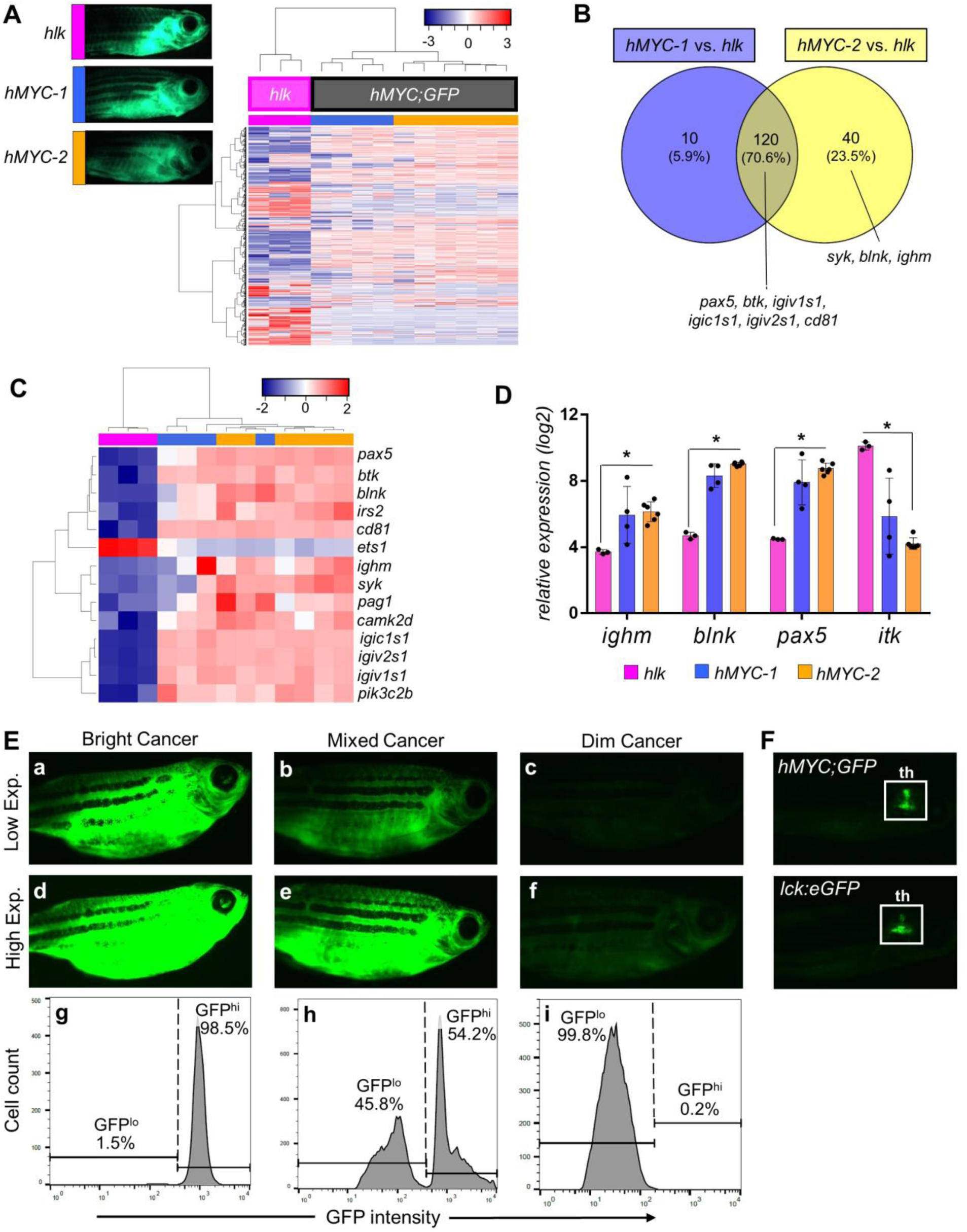
Two ALL types in *hMYC* zebrafish with differing fluorescence intensities. **(A)** Unsupervised analysis of 10 *hMYC* (grey) and 3 *hlk* (magenta) ALL, using highest-variance probes. *hMYC* ALL cluster into *hMYC-1* (blue) and *hMYC-2* (orange) groups. Representative fluorescent images of fish with ALL from each group shown at upper left. **(B)** Venn diagram of 170 over-expressed genes in *hMYC* ALL compared to *hlk* T-ALL. Genes up-regulated by both *hMYC-1* and −2 (n=120) reside in the intersection, including six B cell-specific genes listed below the Venn diagram. Three other B cell-specific genes over-expressed by only *hMYC-2* ALL are listed below the yellow circle. **(C)** Unsupervised analysis using B cell-specific genes. **(D)**Log_2_ expression of *ighm, blnk, pax5* and *itk* in *hlk, hMYC-1* and *hMYC-2* ALL. Each gene is significantly differentially expressed in *hlk* T-ALL versus *hMYC-2* ALL (*p*-value *< 0.05). Results expressed as mean values ± standard deviation (S.D.). **(E) Left:** “bright” ALL, shown using low and high exposure settings (a, d). Cells are GFP^hi^ by flow cytometry (g). **Right:** “dim” ALL, using low and high (c, f) exposures. Cells are GFP^lo^ (i). **Center:** Fish with mixed-ALL (b, e), with discrete GFP^lo^ and GFP^hi^ populations (h). (**F**) Images of control *hMYC;GFP* (upper) and *lck:eGFP* (lower) fish with only thymic (**th**) fluorescence.

Unsupervised analysis divided *hlk* and *hMYC;GFP* malignancies precisely, emphasizing fundamental differences in ALL from different *D. rerio* models (Fig. 1A). Unexpectedly, *hMYC;GFP* ALL also clustered into two subgroups with distinct gene expression profiles (GEPs). To further investigate these groups, we used *hlk* T-ALL as a reference and designated the 4 ALL closest to *hlk* as *hMYC-1,* and the 6 ALL at the far right as *hMYC-2* (blue and orange samples in Fig. 1A).

Separate comparisons of *hMYC-1* or *hMYC-2* vs. *hlk* ALL revealed that B cell-specific genes were up-regulated by both types of *hMYC;GFP* ALL (*pax5, btk, cd81*, etc.; Fig. 1B), with *hMYC-2* ALL over-expressing additional B cell-specific genes (*syk, blnk, ighm*). Ingenuity Pathway Analysis™ (IPA) of these differentially-expressed genes showed enrichment and activation of “*PI3K-Signaling in B-Lymphocytes*”, “*B-cell receptor-signaling*” and *“FcγRIIB-signaling in B-cells*” pathways by *hMYC-2* ALL, but not *hMYC-1*, relative to *hlk* T-ALL (data not shown). To further investigate the unanticipated expression of B cell genes by *hMYC* ALL, we repeated unsupervised analysis using only 14 statistically-significant B cell-specific genes. Remarkably, this signature classified *hlk* vs. *hMYC* ALL perfectly and largely reformed both the *hMYC-1* and *hMYC-2* subclasses (Fig. 1C).

Expression of B cell genes by *hMYC* cancers was unexpected, because B-ALL has never been described by several laboratories—including ours—that study transgenic *Myc*/*MYC* zebrafish (6, 10, 11, 15, 18, 21). Yet microarrays clearly demonstrated B cell genes (*ighm, blnk, pax5*) were expressed at high, medium, and low levels by *hMYC-2, hMYC-1*, and *hlk* ALL, respectively, with T cell-specific *itk* showing the opposite pattern (Fig. 1D). We hypothesized that *hMYC-1* and *-2* cancers might contain different proportions of lymphocytes, with *hlk* being “pure” T-ALL, but *hMYC* ALL comprised of varying amounts of T-ALL, B-ALL, and/or B cells. Alternatively, leukemias can express aberrant markers (22), and *hMYC* might de-differentiate ALL, obscuring cell identities. In either case, B cell genes were high in *hMYC-2* and detectable in *hMYC-1* also, so we next sought to definitively identify the cellular composition of *hMYC* cancers.

### B-ALL and T-ALL each occur in *hMYC;GFP* animals, with different GFP intensities

To definitively identify *hMYC;GFP* ALL as they first developed, we used serial fluorescent microscopy to monitor unaffected animals (i.e., fish lacking visible cancers). In young adults (3-6 months), we observed two phenotypes: brightly-fluorescent cancers originating in thymus and dimly-fluorescent cancers with variable thymic involvement (Figs. 1E, S1). To distinguish these, we used “low-exposure” settings that detected only bright cancers [Figs. 1E(a) vs. (c), S1A vs. S1B], and “high-exposure” settings that revealed dim ALL which were otherwise not visible [Figs. 1E(c) vs. (f), S1B vs. S1D]. Dim ALL differed from non-cancerous *hMYC* and *lck:eGFP* control animals that showed only normal thymic fluorescence (Fig. 1F).

Flow cytometric analyses confirmed microscopy findings, with bright and dim ALL showing distinct, ≥10-fold GFP intensity differences [Fig. 1E(g-i)]. Thus, we could discern ALL with only bright (GFP^hi^), only dim (GFP^lo^), or both cell populations. We analyzed ALL from 27 *hMYC* fish with fluorescent cancers at 6 months of age and found 7 dim ALL with near-exclusively GFP^lo^ cells (Fig. 2A) and 14 GFP^hi^-only ALL (Fig. 2C). Intriguingly, we also found 6 mixed-ALL that contained distinct populations of both GFP^hi^ and GFP^lo^ cells (Fig. 2B). Remarkably, these 27 animals developed 33 total ALL, 13 GFP^lo^ and 20 GFP^hi^.

**Fig. 2.**
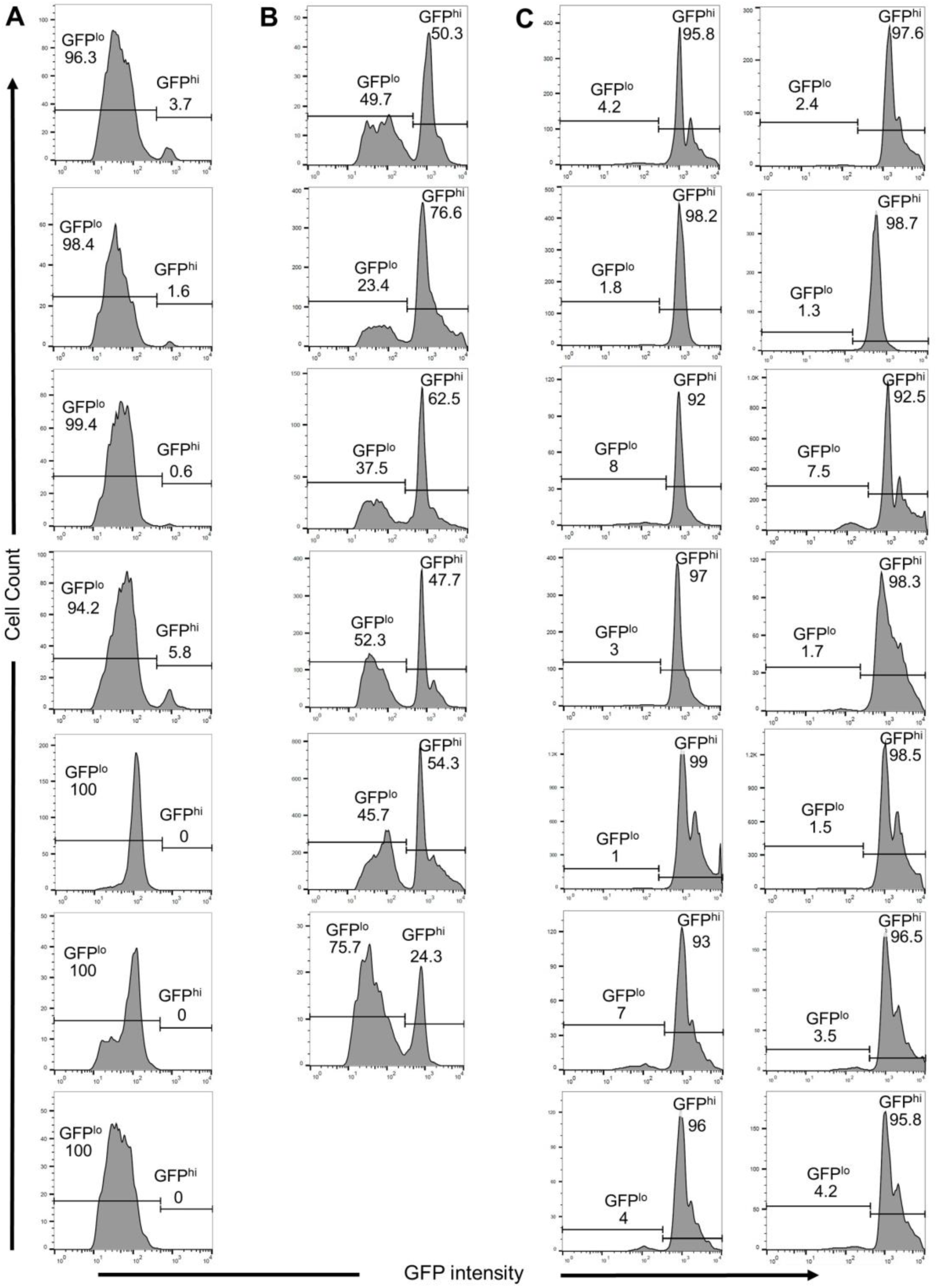
Distinct GFP intensities of *hMYC* dim and bright ALL. Flow cytometric plots of 33 ALL from 27 6-month *hMYC* fish: **(A)** 7 dim, GFP^lo^ ALL **(B)** 6 mixed, GFP^lo^ & GFP^hi^ ALL, and **(C)** 14 bright, GFP^hi^ ALL.

We next tested FACS-purified GFP^+^ dim, bright, or mixed-ALL cells for B cell-(*pax5, cd79b, ighz*, etc.), T cell-(*cd4, cd8, il7r*, etc.), and lymphoblast-(*rag2, igic1s1*, etc.) specific transcripts, as well as the *GFP* and *hMYC* transgenes by quantitative reverse-transcriptase PCR (qRT-PCR), analyzing all GFP^+^ cells as one population without separating GFP^lo^ and GFP^hi^ peaks. Dim/GFP^lo^ ALL expressed B, but not T, cell-specific genes [Fig. 3A(a-d)]. Low *lck* and *GFP* levels in dim ALL matched their weak *in vivo* fluorescence. Conversely, bright/GFP^hi^ cancers expressed only T cell genes. Mixed-ALL expressed both B and T cell genes at intermediate levels. Overall, expression correlated exactly with dim/GFP^lo^ vs. bright/GFP^hi^ phenotypes, and only mixed-ALL, which contained GFP^lo^ and GFP^hi^ cells, co-expressed genes of both cell types. Based on these findings, we conclude dim/GFP^lo^ ALL are B-lineage ALL. Mixed-ALL always exhibited distinct GFP^lo^ and GFP^hi^ cell populations (Fig. 2B) and expressed B-and T-lineage genes [Fig. 3A(a-d)], so we deduce mixed-ALL are not biphenotypic, but simultaneous B-and T-ALL in one animal. B-vs. T-lineage ALL could be unambiguously distinguished by *igic1s1* [Fig. 3A(e)], a homologue of the *IGLL1* surrogate Ig light chain gene expressed by only immature B cells (23). Only dim and mixed-ALL expressed *igic1s1*, but every ALL showed similar levels of the V(D)J recombination enzyme *rag2* [Fig. 3A(f)]. Based on *igic1s1* and *rag2* results, which only immature B cells co-express, we conclude *hMYC* B-ALL are, in fact, pre-B ALL. A zebrafish *rag2* promoter regulates *hMYC*, so it is logical that *rag2^+^* B-lymphoblasts (i.e., pre-B cells) are affected, just like T-ALL in this model (6, 10). Moreover, similar *hMYC* levels in pre-B and T-ALL [Fig. 3A(f)] indicate this transgene has similar oncogenic potency in both lymphocyte lineages. Surprisingly, *rag1* was much lower in pre-B ALL [Fig. 3A(e)], making *igic1s1* and *rag1* a two-gene panel that can distinguish *hMYC* ALL types independent of *lck* or *GFP* levels. Mammalian B-and T-lymphoblasts co-express RAG1 and RAG2, so dichotomous *rag1* levels were unexpected. However, unlike mammals, *rag2*-mutant zebrafish lack T cells, yet retain functional B cells (24), further suggesting V(D)J recombination may differ between mammals and *D. rerio*.

**Fig. 3.**
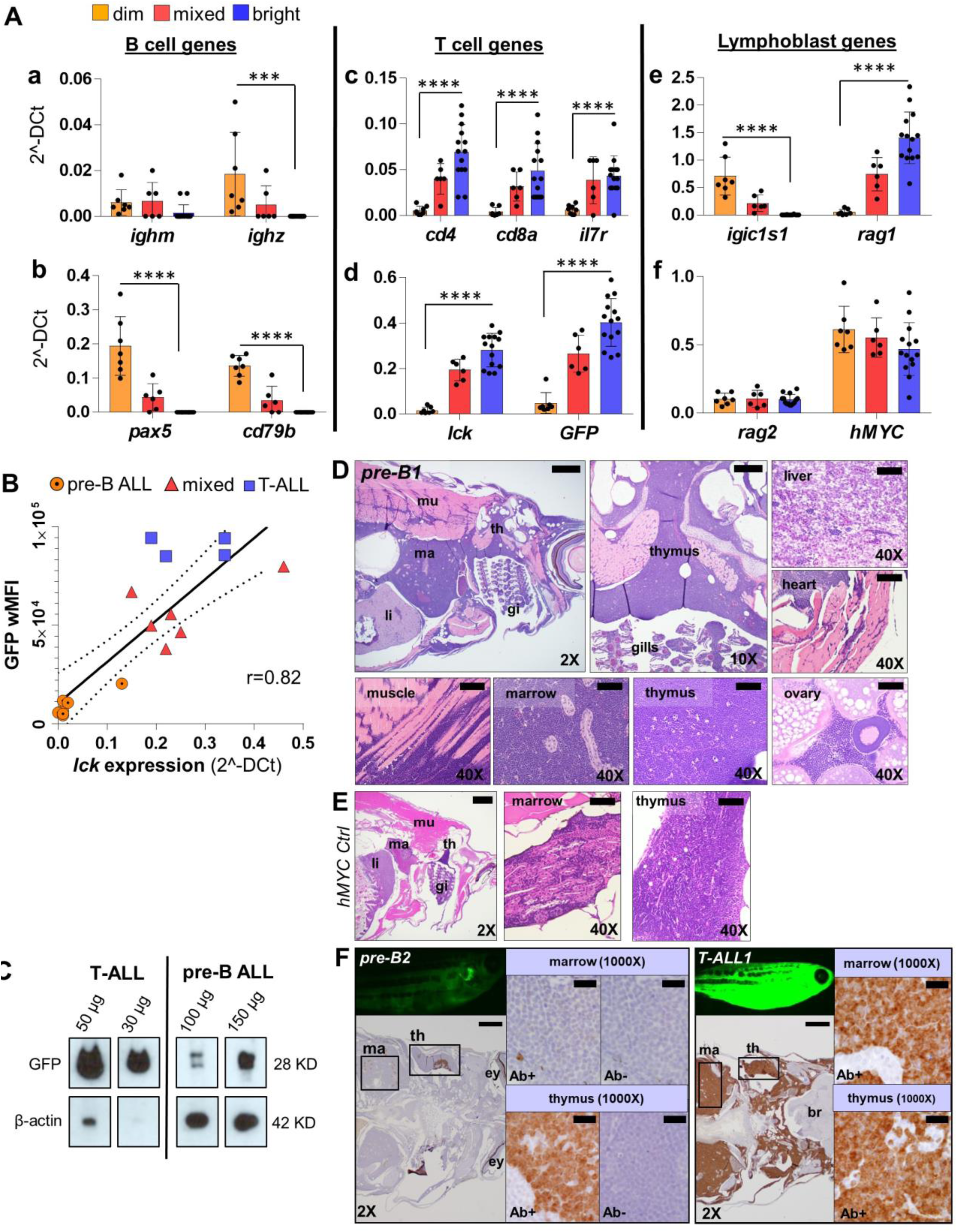
GFP fluorescence intensity of *hMYC* ALL correlates with B vs. T cell lineage. **(A)** qRT-PCR of ALL with differing GFP fluorescence (dim, n=7; mixed, n=6; bright, n=14) of B cell-specific (*ighm, ighz, pax5, cd79b*; a-b), T cell-specific (*cd4, cd8a, il7r, lck*; c-d), lymphoblast-specific (*igic1s1, rag1, rag2*; e-f) genes and transgenes (*GFP,* d; *hMYC*, f). Results shown as mean values ± S.D., after normalization to housekeeping genes (*β-actin* and *eef1a1l1*). Significant differences are indicated (*p-values*: ***<0.001, ****<0.0001). (**B**) Spearman correlation (*p*-value 0.0002, r = 0.82, r^2^ = 0.74) between *lck* vs. wMFI for: 7 B-ALL (circles), 6 mixed (triangles), and 4 T-ALL (squares). The solid line represents linear regression; dashed lines denote 95% confidence intervals. (**C**) Anti-GFP and anti-β-actin WB of FACS-purified T-and pre-B ALL. Note different amounts of total protein loaded. (**D**) H&E of pre-B ALL infiltration in different sites and (**E**) *hMYC* control. (**F**) Anti-GFP IHC of *hMYC* pre-B ALL (*pre-B2*, left) and T-ALL (*T-ALL1*, right). 1000X images show staining with or without anti-GFP (Ab+; Ab-). Abbreviations: **th**=thymus, **ma**=marrow, **li**=liver, **mu**=muscle, **gi**=gills, **ey**=eye, **br**=brain. 2X scale bar=500 µm; 10X bar=100 µm; 40X bar=50 µm; 1000X bar=20 µm.

As predicted by different *in vivo* fluorescence [Figs. 1E(a-f), S1] and GFP^lo^ vs. GFP^hi^ cytometric results [Figs. 1E(g-i), 2], *lck* and *GFP* also differed markedly between pre-B and T-ALL [Fig. 3A(d)]. A zebrafish *lck* promoter regulates GFP (20), and pre-B ALL expressed little *lck* or *GFP*, while T-ALL expressed both abundantly. In agreement, *lck* mRNA correlated with weighted median fluorescence intensity (wMFI; Fig. 3B), with each ALL type clustering separately, proving *lck* levels—and thus, cellular fluorescence—distinguish pre-B vs. T-ALL in this model. We also confirmed *GFP* mRNA and protein levels agree by Western blot (WB), with much higher amounts of total protein needed to detect GFP in pre-B ALL compared to T-ALL (Fig. 3C).

*lck* is generally considered to be T cell-specific (20), but zebrafish NK and myeloid cells also express *lck* (25, 26). Pertinent to our study, we analyzed public data from different maturation stages of human lymphocytes (27) and found pre-B cells expressed higher *LCK* than naïve and mature B cells, although below that of T cells (Fig. S2A). Additionally, Microarray Innovations in Leukemia (MILE) data (28) from 1,816 leukemia patients also showed *LCK* levels in pre-B ALL equal to many T-ALL (Fig. S2B). Thus, human and zebrafish pre-B ALL both express *LCK*/*lck*.

### Zebrafish pre-B ALL resembles human pre-B ALL morphologically

To examine pre-B ALL histology, we analyzed *hMYC* fish with dim ALL. Hematoxylin and eosin (H&E) stains showed lymphoblast infiltration of the kidney-marrow, thymus, liver, and elsewhere (Figs. 3D, S3A). Marrow hypertrophy was often profound, with marrow expansion and invasion through muscle into subcutaneous tissue and skin (Fig. S3A, *pre-B3, -B4*). Yet despite these massive disease burdens, fish remained only dimly fluorescent. As in humans, pre-B and T-ALL were indistinguishable by H&E (Fig. S3A-B), and both were markedly abnormal compared to control animals (Figs. 3E, S3C).

Immunohistochemical analysis (IHC), however, could discriminate pre-B from T-ALL, with very faint anti-GFP staining in GFP^lo^ ALL vs. strong signals in GFP^hi^ ALL (Figs. 3F, *pre-B2* vs. *T-ALL1*, S4A-B), and only remnant thymic tissue showing strong signals in pre-B ALL fish (Figs. 3F, *pre-B2* and S4A, *pre-B5, -B6*). Consistent with this, regions stained weakly by anti-GFP (Ab+) corresponded to dimly-fluorescent anatomic regions (Figs. 3F, S4A; 1000X panels). Because Ab recognizing zebrafish lymphocyte proteins are not available, we used RNA *in situ* hybridization (ISH; RNAscope™) to independently test cell identities using probes for *hMYC* and B cell-specific *cd79b* (Fig. 4A-C). *hMYC* labeled pre-B ALL strongly (Fig. 4A; H&E of this animal shown in Fig. 3D), including cells in the thymus and kidney-marrow (Fig. 4B-C, *pre-B1*). These same areas were also *cd79b*-positive, confirming B-lineage. Thymi of *hMYC* control fish were avidly *hMYC*^+^, but had few *cd79b^+^* cells (Fig. 4B, D, *hMYC Ctrl*), indicating thymic B cells are sparse unless pre-B ALL is present. Similarly, *hMYC* control marrow had fewer dually *hMYC*^+^/*cd79b^+^* cells (Fig. 4C-D), with normal kidney-marrow architecture, including renal tubules. *hMYC* was absent in the thymus of control *lck:eGFP* fish (Fig. 4B, *lck:eGFP*) proving probe specificity, and showed rare *cd79b^+^* B cells, demonstrating few thymic B cells in WT fish. We examined a different animal with disseminated pre-BALL and localized T-ALL, based on microscopy and IHC findings (see microscopy, IHC, and H&E in Fig. S4A, *pre-B6*), adding a T-cell specific probe, *lat*, to distinguish pre-B vs. T-ALL. RNA ISH demonstrated cells that were *hMYC*^+^/*cd79b*^+^/*lat*^-^had completely replaced the marrow and thymic cortex (Fig. 4E), with GFP^hi^ *hMYC*^+^/*cd79b*^-^/*lat*^+^ cells remaining only in an enlarged thymic medulla (i.e., localized T-ALL). Similar results were seen in a second animal with near-complete thymic ablation by pre-B ALL (Fig. S4C; Fig. 3F, *pre-B2* shows microscopy and IHC of this specimen). In summary, *in vivo* fluorescence, cytometric GFP intensity, qRT-PCR, WB, and RNA ISH all prove dim cancers in *hMYC*;*GFP* fish are pre-B ALL with organ distributions similar to human pre-B ALL.

**Fig. 4.**
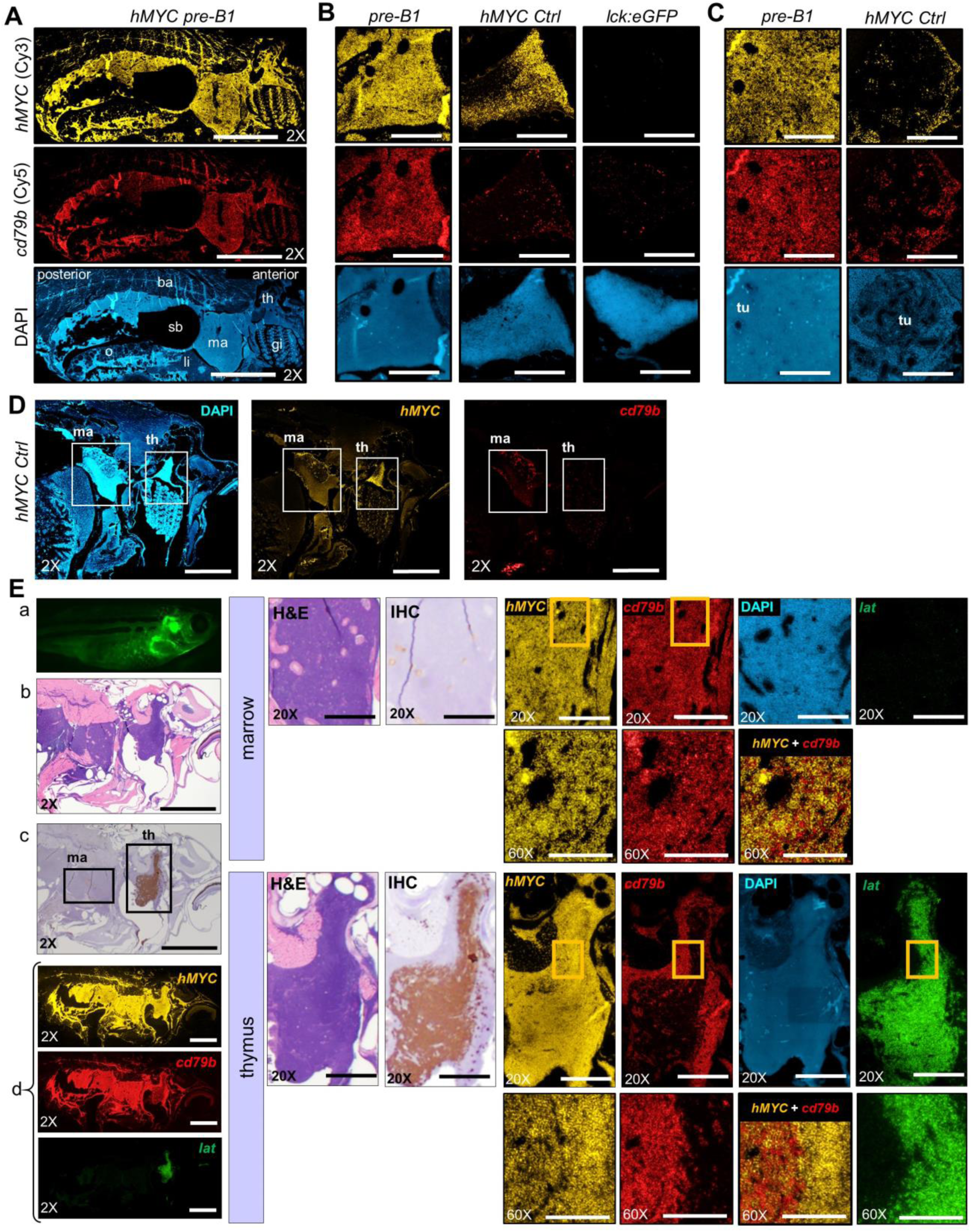
Pre-B ALL co-express *hMYC* and B cell-specific *cd79b*. RNA ISH for *hMYC* (yellow) and *cd79b* (red) in: (**A**) sagittally-sectioned *hMYC* pre-B ALL (*pre-B1*; scale bar=2 mm), (**B**) Thymi of *pre-B1* (left), *hMYC* control (center), and *lck:GFP* (WT) control (right; scale bar=200 μm), (**C**) Kidney-marrow of *pre-B1* (left) and *hMYC* control (right). Kidney tubules (**tu**) are displaced by *pre-B1* ALL cells in marrow DAPI image (scale bar=200 μm), (**D**) *hMYC* control (scale bar=1 mm). (**E**) Second *hMYC* fish with pre-B ALL and localized thymic T-ALL. **Left:** (a) High-exposure microscopy, (b) H&E, (c) Anti-GFP IHC, (d) RNA ISH for *hMYC* (yellow), *cd79b* (red) and *lat* (green). **Middle:** high-power of kidney-marrow (top) and thymus (bottom) by H&E, anti-GFP IHC. **Right:** *hMYC, cd79b*, and *lat*, RNA ISH. Boxed regions in 20X marrow and thymus panels are enlarged in the 60X images directly beneath them. Merged *hMYC*+*cd79b* images are also shown. 2X scale bar=1 mm; 20X bar=200 µm; 60X bar= 100 µm. Abbreviations as in Fig. 3, and **o**=ovary; **ba**=back; **sb**=swim bladder.

### Zebrafish pre-B ALL remain GFP^lo^/*lck*^lo^ in every tissue

Pre-B ALL disseminated aggressively (Figs. 3D-F, 4, S3A, S4A, C). To test whether pre-B ALL cells retained a GFP^lo^/*lck*^lo^ phenotype in every niche, we examined thymus, marrow, spleen, blood, and muscle and viscera by flow cytometry (Fig. 5). Fish with pre-B ALL (n=5, B1-5) showed GFP^lo^ cells in each tissue, with GFP^hi^ cells (i.e., T cells) present only in thymus. Two fish (B1, B2) had numerous thymic GFP^lo^ cells, demonstrating extensive invasion. *hMYC* T-ALL controls (n=3, T1-3) exhibited near-exclusively GFP^hi^ populations in every tissue. We also analyzed one mixed-ALL (M1). Consistent with two co-existing ALLs, the thymus, spleen, and muscle and viscera all contained large proportions of both GFP^lo^ and GFP^hi^ cells, with GFP^lo^ cells outnumbering T-ALL cells in marrow and peripheral blood. To conclusively test whether GFP^lo^ and GFP^hi^ cells always equate to B-vs. T-lineage, we analyzed *igic1s1* and *rag1* in FACS-purified GFP^lo^ and GFP^hi^ cells from 5 of these fish (B1, B2, M1, T1, T2). As previously [Fig. 3A(e)], dim cells from every tissue expressed only *igic1s1*, and only bright cells were *rag1*^+^, including the GFP^lo^ and GFP^hi^ ALLs of M1. Thus, GFP reliably reflects pre-B vs. T-ALL in any niche, establishing *hMYC*;*GFP* zebrafish as a new and novel model to study both ALL types in one genetic context, or even one animal.

**Fig. 5.**
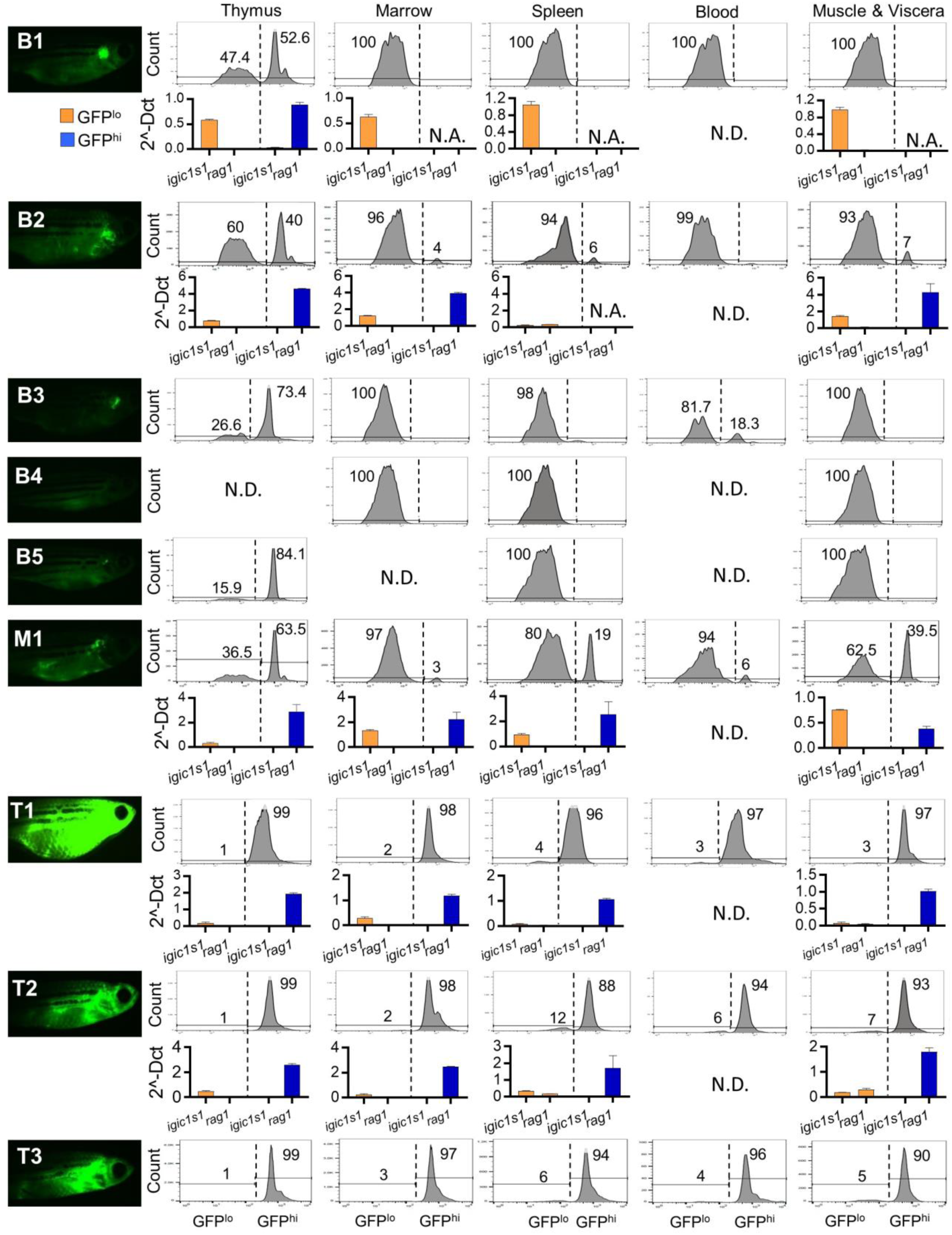
GFP intensity and *igic1s1*/*rag1* distinguish pre-B vs. T-ALL in each anatomic site. Left column shows high-exposure fluorescent microscopy of 4 month-fish with pre-B ALL (B1-5), mixed ALL (M1), or T-ALL (T1-3). Panels at right show flow cytometric analysis of GFP^lo^ and GFP^hi^ cells of thymus, marrow, spleen, peripheral blood, and muscle & abdominal viscera. Each panel shows % of GFP^lo^ vs. GFP^hi^ cells in the entire GFP^+^ gate; 10^5^ events from the lymphoid/precursor gate were analyzed for each plot. N.D. = not determined. Histograms depict expression of *igic1s1* and *rag1* by qRT-PCR in five *hMYC* fish with pre-B ALL (B1, B2), mixed ALL (M1) or T-ALL (T1, T2). Results shown as mean ± S.D. with normalization relative to *β-actin* and *eef1a1l1* housekeeping genes. N.A. = not available due to insufficient cells for RNA extraction.

### MYC-induced zebrafish pre-B and T-ALL have distinct expression signatures

To test whether *D. rerio* and human pre-B ALL share similar gene expression, we next defined GEPs in a new cohort of animals, quantifying 96 transcripts that distinguish B/T/NK cells, lymphoblasts, and precursor populations (genelist in Table S1) (29, 30). We FACS-purified 8 pre-B ALL, 4 T-ALL, and 2 ALL from a mixed-ALL fish (Fig. 6A-C; GFP^lo^ or GFP^hi^ populations in orange or blue, respectively), as well as control lymphocytes. For T cell controls, we isolated GFP^hi^ thymocytes from three 10-fish cohorts of control *hMYC* and *lck*:*GFP* fish (*hMYC* thymus-GFP^hi^, WT thymus-GFP^hi^; Fig. 6D). As another T cell control, we pooled marrow, lymphoid-gated (31) GFP^hi^ cells from these same 30 WT fish (WT marrow-GFP^hi^; Fig. 6E, blue). *hMYC* marrow lacked GFP^hi^ cells (Fig. 6F), so these were not analyzed. For B cells, we purified GFP^lo^ and GFP^-^ marrow cells of the same 3 *hMYC* control cohorts (*hMYC* marrow-GFP^lo^, *hMYC* marrow-GFP^-^; Fig. 6F) and from marrow of 30 WT fish used for thymocyte preparations (WT marrow-GFP^lo^, WT marrow-GFP^-^; Fig. 6E).

**Fig. 6.**
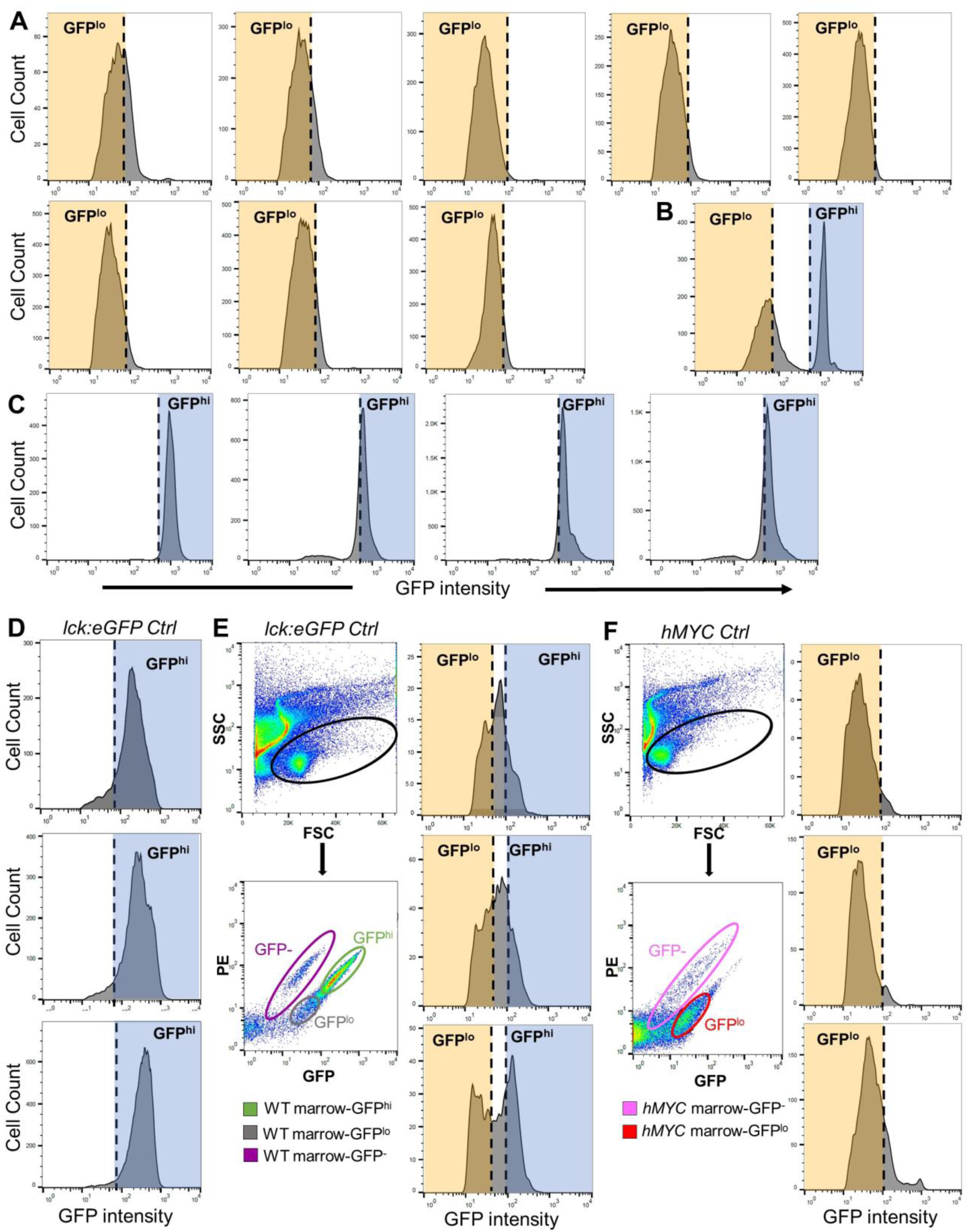
GFP^lo^ and GFP^hi^ lymphocytes isolated for expression profiling. Plots of GFP^lo^ (orange) and GFP^hi^ (blue) populations FACS-purified for RNA quantification. (**A**) Pre-B ALL (n=8) sorted as pure GFP^lo^ populations (orange); (**B**) Mixed ALL sorted as separate GFP^lo^ (orange) and GFP^hi^ (blue) populations; (**C**) T-ALL (n=4) sorted as pure GFP^hi^ populations (blue). Control lymphocyte populations: (**D**) 3 groups of pooled WT *lck:eGFP* thymi (each group, n=10 fish) sorted for GFP^hi^ thymocytes (blue), (**E-F**) 3 groups of pooled WT or *hMYC* marrow (each group, n=10 fish). Upper left panels show side-and forward-scatter (SSC, FSC) gating of the lymphocyte/precursor population (black ovals). Panels below show subsequent purifications of these cells into GFP^-^ (fuchsia and pink ovals), GFP^lo^ (gray and red ovals) and GFP^hi^ (green oval) sub-fractions. Note paucity of GFP^hi^ cells in *hMYC* marrows, which were not isolated. Oval colors match color-coding in Fig. 7. Triplicate plots at right show GFP^lo^ and GFP^hi^ biologic replicates for each 10-fish marrow sample. Marrow cells from each WT group (GFP^-^, GFP^lo^, GFP^hi^) were pooled from all 30 fish to yield sufficient RNA for expression analysis.

Using barcoded gene-specific probes (Nanostring nCounter™), we quantified mRNA levels of 96 transcripts in all 29 samples. Unsupervised analysis clustered all B and T cell triplicate controls tightly (*hMYC* thymus-GFP^hi^, WT thymus-GFP^hi^, *hMYC* marrow-GFP^-^, *hMYC* marrow-GFP^lo^), proving high reproducibility of biologic replicates (Fig. 7A). Each of the 29 samples segregated unequivocally as B-or T-lineage, with every GFP^-^/GFP^lo^ sample grouping as B-lineage (n=17), and GFP^hi^ samples forming a T-lineage group (n=12). Every GEP clustered perfectly according to GFP expression, whether from control fish or fish with ALL, and irrespective of whether cells were obtained from thymus or marrow. Notably, GFP^-^ and GFP^lo^ cells from WT and *hMYC* animals all exhibited similar B-lineage profiles, with *hMYC* GFP^lo^ B cell GEPs most similar to pre-B ALL (Fig. 7A).

**Fig. 7.**
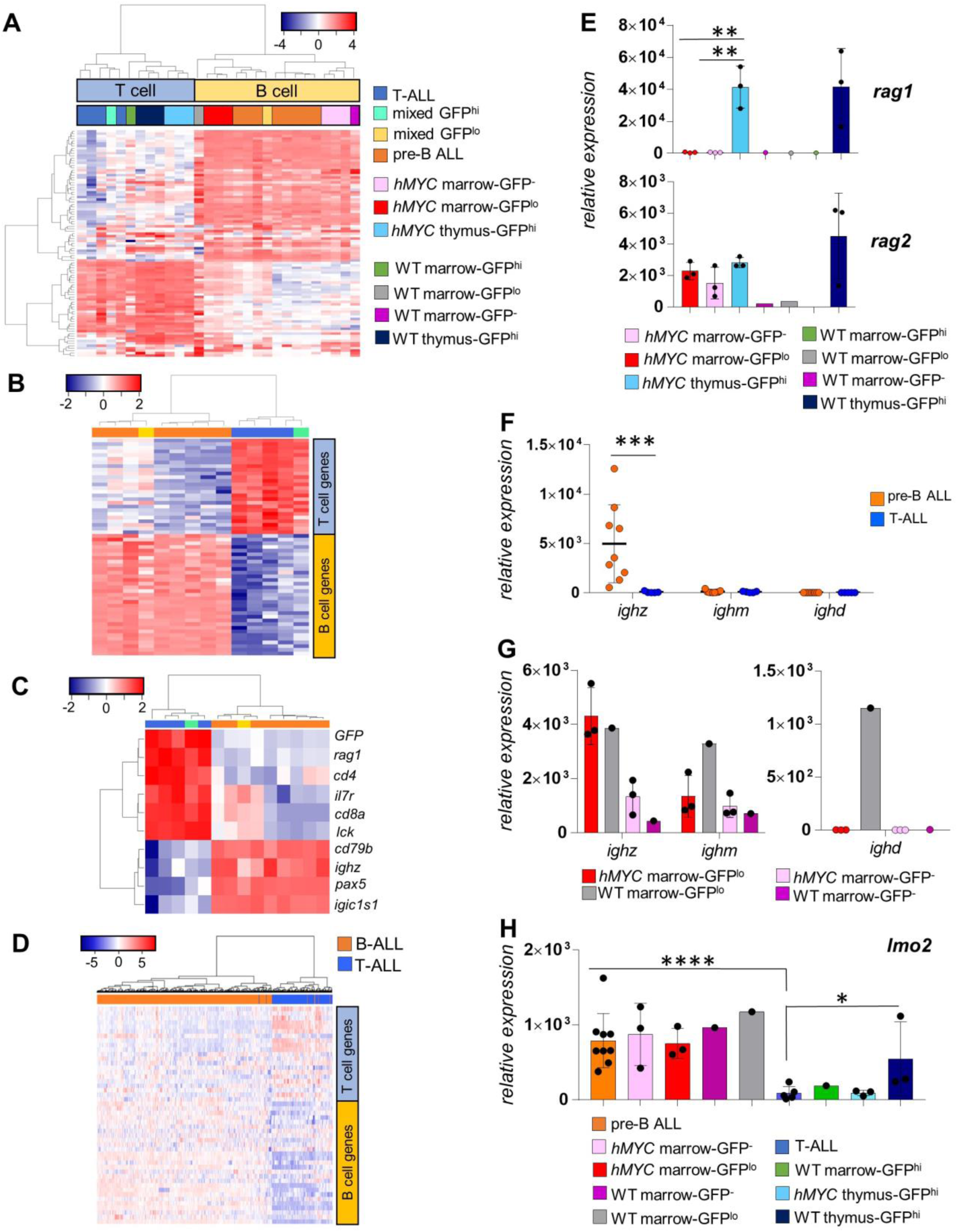
*hMYC* drives pre-B and T-ALL with distinct GEP and alters B-lineage expression. (**A**) Unsupervised analysis of all malignant and normal lymphocyte populations (n=29) based on GEPs of 93 genes (*β-actin, eef1a1l1, gapdh* housekeeping genes used for normalization not shown). Each sample groups as T-(blue box at top; n=12) or B-lineage (yellow box; n=17). Gene order listed in column D of Table S1. **(B)** Supervised analysis using significant (FDR<0.05) genes (n=59; order in Table S2) distinguishing pre-B vs. T-ALL. Pre-B (orange; n=8), T-(blue; n=4), and two ALL from a mixed-ALL (GFP^lo^=yellow, GFP^hi^ =green) cluster as pre-B or T-ALL. (**C**) Analysis with genes from prior qRT-PCR testing (Fig. 3A). (**D**) Unsupervised clustering of human ALL from patients on the MILE1 study, using homologues of zebrafish genes that distinguish *hMYC* pre-B vs. T-ALL (human genes in same order as Table S2). (**E**) *rag1* (top) and *rag2* (bottom) expression in *hMYC* and WT control B and T cell populations. (**F**) *ighz, ighm* and *ighd* levels in pre-B (orange) and T-ALL (blue). (**G**) *ighz, ighm* and *ighd* expression in GFP^lo^ (red) or GFP^-^ (pink) non-malignant *hMYC* B cells, and GFP^lo^ (grey) or GFP^-^ (fuchsia) WT B cells. (**H**) *lmo2* levels in all samples (n=29) showing highest expression in B-lineage groups and lowest expression in T-ALL. Mean values are shown ± S.D., with significant differences noted (*p-values*: *<0.01, **<0.001, ***<0.001, ****<0.0001).

Pre-B and T-ALL GEPs were distinct (Fig. 7B; Table S2 lists differentially-expressed genes), with the GFP^hi^ ALL (green) and GFP^lo^ ALL (yellow) of the mixed-ALL animal grouping as T-or pre-B ALL, respectively. In total, ∼60 homologous genes able to distinguish human (30) and zebrafish (29) B vs. T cells likewise categorized *hMYC* pre-B vs. T-ALL. Key classifier-genes (Fig. 7C) matched prior qRT-PCR results (Fig. 3A), with both ALL types showing comparable levels of *rag2, hMYC*, and *shmt2*, a known direct MYC target (Fig. S5A) (21), reinforcing that *hMYC* levels and activity are similar in this dual pre-B/T-ALL model. To further examine conservation of gene expression between both types of *D. rerio* and human ALL, we also tested whether human homologues of the differentially-expressed *hMYC* pre-B-vs. T-ALL genes could reliably classify ALL from the MILE1 study (Fig. 7D). This signature exhibited remarkable classification power, separating nearly all of these human ALL correctly. Homologues of several other hematopoietic stem/progenitor-or immature lymphocyte-specific genes showed no significant difference between *hMYC* ALL types (Fig. S5B).

Like ALL, control *hMYC* and WT thymocytes expressed more *rag1* than B cell controls (Fig. 7E, top). From this result, we deduce higher *rag1* in zebrafish T-ALL vs. pre-B ALL is unrelated to malignancy, but a normal feature of *D. rerio* lymphoblasts, although different from mammals. Notably, *hMYC* B cell controls showed higher *rag2* than WT B cells, implying *hMYC* may expand the pre-B cell population (Fig. 7E, bottom). We also analyzed Ig heavy chain expression. Surprisingly, pre-B ALL expressed only *ighz* (Fig. 7F), an isotype unique to teleost fish that is functionally analogous to mammalian IgA (32). GFP^lo^ *hMYC* B cells expressed *ighm* (Fig. 7G, left), so *ighm*^+^ pre-B ALL should be detectable in *hMYC* fish—but these were not found. Therefore, MYC may be oncogenic in only the *ighz*-lineage. Another B-lineage abnormality we noted is that *hMYC* GFP^lo^ cells lacked *ighd* (Fig. 7G, right), so *hMYC* may suppress *ighd* transcription or repress this lineage, just as it seems to block the T cell lineage in *hMYC* marrow (Fig. 6F). Overall, *hMYC* dramatically perturbs zebrafish B cell development, inducing ALL in *ighz*^+^ B cells, and skewing both the *ighm* and *ighd* lineages.

Finally, we found substantially higher *lmo2* in normal and malignant B vs. T cells (Fig. 7H), mirroring *LMO2* findings in human B vs. T cells (33), which we independently confirmed in public human B and T cell data (Fig. S5C). This contradicts the assertion that MYC-driven zebrafish T-ALL emulates the human TAL1/LMO subtype (6). Instead, *hMYC* T-ALL actually had the lowest *lmo2* levels of the 9 populations that were tested, including all T cell controls (Fig. 7H).

## Discussion

Pre-B ALL is the most common pediatric cancer and kills more children than any other type (3), but no good zebrafish models exist for this important disease. Drug screens (17, 18), genetic studies (11, 12, 15) and stem cell discoveries (13, 14) were all made possible by zebrafish T-ALL models, and these and other approaches would likely be similarly fruitful with *D. rerio* pre-B ALL. Here, we describe the first robust zebrafish pre-B ALL model. Unexpectedly, young 3-6 month *hMYC* fish—used for years in several of the aforementioned T-ALL studies—also develop highly-penetrant pre-B ALL. Remarkably, this went unrecognized for over a decade. In terms of detecting pre-B ALL, dual-transgenic *hMYC*;*GFP* fish proved particularly valuable to our study, because their *lck:GFP* expression not only allowed pre-B ALL to be detected, but their differing GFP levels also distinguished pre-B and T-ALL *in vivo*. This dichotomy in *lck* expression extends to normal B and T cells as well, is corroborated by flow cytometry, and corresponds precisely to B-and T-lineage GEPs (Figs. 1E, 2-7, S1, S4). Consequently, even in fish with concomitant pre-B and T-ALL—which we believe are unique to this model—these cell lineages and ALL types, can be reliably separated for independent study.

Apart from the utility of *lck:GFP* in this system, *hMYC* pre-B ALL are powerful because they emulate this human disease in several ways: histology and organ involvement (Figs. 3-4, S3-4), *lck* levels (Fig. S2) and, most importantly, gene expression signature (Fig. 7). In fact, genes that differentiate zebrafish pre-B vs. T-ALL (Fig. 7B) classify human ALL also (Fig. 7D). The GEPs we obtained revealed many transcripts that distinguish pre-B vs. T-ALL in this model, but we also report a two-gene panel to categorize *hMYC* ALL that requires only *igic1s1* and *rag1*, and this panel can be applied to any *hMYC* genetic background.

Interestingly, although *RAG1* is expressed by mammalian B-and T-lymphoblasts, we find that only *D. rerio* T-lymphoblasts express *rag1* highly, with levels >70-and >155-fold greater in normal T or T-ALL cells than in B or pre-B ALL cells, respectively (Figs. 3A(e), 5, 7C,E). A recent report of zebrafish B cell development despite low *rag1* supports our observation (34). Apart from *rag1*, pre-B ALL expressed other classic B-lymphoblast genes like *rag2*, surrogate light chain components, *pax5, cd79a/b,* and others (Figs. 3-4, 7C, S4C, S5A). This was true for every dim ALL, including GFP^lo^ mixed-ALL, so we conclude mixed-ALL are co-existing pre-B and T-ALL, and not biphenotypic ALL. Certainly, because *hMYC* can induce both ALL types, it remains possible that mixed-lineage biphenotypic ALL may arise in this system, but to date, our analyses of >40 dim ALL have failed to detect any that express T cell genes, suggesting that GFP^lo^ ALL are always B-lineage, and GFP^hi^ always represent T-ALL. In future work, studying both ALL types in one background—or one animal—presents new opportunities, like efforts to find cooperating genetic lesions unique to one type of ALL, lymphocyte lineage-specific drugs, or myriad other applications.

Our results also demonstrate multiple features of abnormal B and T lymphopoiesis in *hMYC* fish. Of interest, pre-B ALL GEPs closely matched the gene expression pattern of a recently-described *ighz*^+^ B cell population dubbed “fraction 2” (34), suggesting *hMYC* is oncogenically active in this cell population. Supporting this, every pre-B ALL we identified was *ighz*^+^ (Fig. 7F). Liu et al. postulated that the zebrafish *ighm*-lineage lacks a classic pre-B stage. If this is correct, it is logical that pre-B ALL only occurs in *ighz*^+^ cells, the lineage that has pre-B cells. We note that *ighm*^+^ cells do express *lck* (Fig. 7G), so *ighm*^+^ pre-B ALL should be GFP^+^ and detectable in this system. Yet *ighm*^+^ pre-B ALL were never detected, so we conclude they do not occur. In addition to our finding that ALL develops in only *ighz*^+^ cells, we also found that *hMYC* alters other B-lineages, with *ighm* and *ighd* both reduced in *hMYC* marrow (Fig 7G). Whether this is due to *ighm* and *ighd* transcriptional—or lineage—repression remains to be determined. Notably, T cells are also diminished in *hMYC* marrow (Fig. 6F), so *hMYC* alters both the B and T lymphocyte lineages, beyond inducing both pre-B and T-ALL.

Clearly, *hMYC* is leukemogenic to lymphocytes and perturbs zebrafish lymphocyte development. In future work, *hMYC* pre-B ALL can be used for classic zebrafish approaches like chemical and genetic screens, or in mechanistic studies probing *hMYC* biology in either ALL type. *MYC* is arguably the most clinically-relevant oncogene, important in many cancers besides ALL (35), but MYC’s contrasting actions in distinct neoplasias remain largely unexplored. We show *hMYC* fish provide a novel system to address this topic via studies of both human ALL types, using a single model.

## Materials and Methods

### Zebrafish Care and Fluorescent Microscopy Screening

Zebrafish housed in an aquatic colony at 28.5°C on a 14:10 hour light:dark circadian cycle and cared for according to protocols approved by the University of Oklahoma Health Sciences Center IACUC (12-066 and 15-046). For all procedures, fish were anesthetized with 0.02% tricaine methanesulfonate (MS-222). 3-6 month old *hMYC;GFP* fish of both genders were screened for abnormal GFP patterns, using a Nikon AZ100 fluorescent microscope. Low exposure (200 ms, 2.8X gain) and high exposure (1.5s, 3.4X gain) settings were used to obtain images with Nikon DS-Qi1MC camera. Images were processed with Nikon NIS Elements Version 4.13 software.

### Fluorescence-Activated Cell Sorting (FACS) and Flow Cytometry Analysis

Cells from whole fish, excluding head regions, or specific organs were dissociated using a pestle and passed twice through 35 μm filters. GFP^+^ cells were collected from the lymphoid and precursor gates (31) using a BD-FACSJazz Instrument. GFP intensity defined characteristic peaks on either side of ∼10^2^ on the GFP intensity axis of this instrument; this value was used to discriminate between GFP^lo^ and GFP^hi^ peaks. For percentage calculations, all GFP^+^ cells from 10^5^ total events in the lymphoid/precursor gate were evaluated. GFP weighted mean fluorescence intensity (wMFI) was calculated using the following formula: [(MFI of GFP^lo^ population x % of GFP^lo^ cells) + (MFI of GFP^hi^ population x % of GFP^hi^ cells)]. Flow cytometric analyses performed using FlowJo software.

### RNA extraction and quantitative real-time polymerase chain reactions

Total RNA was extracted using Trizol according to manufacturer instructions (Invitrogen). For qRT-PCR, 16 ng of total RNA was reverse transcribed using standard methodology. SYBR-Green qRT-PCR was performed using a CFX96 Touch^TM^ Real-Time PCR Detection System (Biorad). Each cDNA sample was tested in triplicate. The 2^-DeltaCt^ (Delta Ct = Ct_experimental gene_-Ct_housekeeping gene_) method (36) was used to calculate the relative expression of each gene to housekeeping genes (*β-actin* and *eef1a1l1*).

### RNA Microarrays

*In vitro* transcription, hybridization to Zebrafish Genome Arrays (#900487), and biotin-labeling performed according to manufacturer instructions (Affymetrix). Microarray (.CEL file) data generated using Affymetrix GeneChip Command Console Software, normalized against the entire dataset using the justRMA algorithm, and analyzed using R-Bioconductor (Version 3.4.1). Unsupervised and supervised hierarchical clustering were used to group specimens based on Euclidean distance and the Ward method. Differentially-expressed genes identified by shrinkage t-tests (37) with local false discovery rate (lfdr) used to correct *p*-values. A lfdr <0.05 was considered significant for differentially-expressed probe-sets. Public microarray data from normal human B and T cells at various maturational stages (27) and leukemia patients on the MILE stage 1 study (28) were used for gene expression analysis.

### Histology and Immunohistochemistry (IHC)

H&E stains performed using standard methodology. IHC stains performed using 1:8000 dilutions of anti-GFP antibody (GeneTex, #GTX20290) at 37°C using a Ventana BenchMark XT instrument. Staining of GFP^+^ tissues performed using an Inview 3,3’-Diaminobenzidine (DAB) detection kit (Ventana Medical Systems).

### Western Blot Analysis

FACS-purified GFP^lo^ pre-B ALL and GFP^hi^ T-ALL cells were homogenized in lysis buffer [(20mM Tris-HCl pH 8.0, 137mM NaCl, 2mM EDTA, 1% NP-40, and 10% Glycerol supplemented with Protease Inhibitor Cocktail (Sigma, #P8340)]. Total protein was resolved on a 4„10% gradient polyacrylamide gel and transferred to a nitrocellulose membrane (both BioRad) using transfer buffer (25mM Tris, 192mM glycine, and 20% methanol). Tris-buffered saline, 5% non-fat dry milk, and 0.05% Tween-20, used for blocking. Blots incubated overnight at 4°C with anti-GFP primary antibody (Santa Cruz, #sc-9996), followed by incubation with horseradish peroxidase-conjugated secondary antibody (BD). Immuno-reactive bands detected using ECL 2 western blot substrate (Pierce). Parallel incubation with anti-β-actin antibody (Abcam, #ab8227) used as a positive control.

### RNAscope-ultrasensitive in situ hybridization (RNA ISH)

RNAscope (Advanced Cell Diagnostics-ACD) fluorescent-field ISH was used to detect *hMYC, cd79b* and *lat* mRNA in fish sections. Fixation, sectioning, and staining performed using the RNAscope Multiplex Fluorescent Detection Kit v2 (#323110), according to manufacturer instructions (https://acdbio.com/). RNAscope probes (ACD) used to specifically detect *human MYC* (#311761-C2), *D. rerio cd79b* (#511481) and *lat* (#507681). Probe labels (PerkinElmer) as follows: TSA-Plus-Cyanine-3 (#NEL744001KT) for *hMYC* (yellow fluorescence), TSA-Plus-Cyanine-5 (#NEL745001KT) for *cd79b* (red), and TSA-Fluorescein (#NEL701A001KT) for *lat* (green). Slides imaged and analyzed using Operetta High-Content Imaging System (PerkinElmer) and Harmony 4.1 software.

### Nanostring nCounter Gene Expression Profiling

GEPs of FACS-purified GFP^lo^ and GFP^hi^ cell populations were quantified using a 96-gene Custom CodeSet according to manufacturer instructions (Nanostring nCounter Technologies). Genes quantified using an nCounter Digital Analyzer and analysed using nSolver v3.0 software. Background thresholds defined by counts from a no-RNA blank that were subtracted from each sample. Raw counts were normalized to spiked-in positive control probes and housekeeping genes (*β-actin, eef1a1l1* and *gapdh*), as suggested by the manufacturer. nSolver t-tests used to compare groups and identify differentially-expressed genes (FDR ≤0.05).

### Statistical analysis

GraphPad Prism 7 software used to calculate Spearman correlations and non-parametric Mann-Whitney t-tests for genes tested by qRT-PCR. Two-tailed 95% confidence intervals were used to determine significance, and significant differences are reported as *p-values* with *p\** <0.05, ** <0.01, *** <0.001 and **** <0.0001.

## Acknowledgments

We thank Megan-Malone Perez at OUHSC for zebrafish care, Sheryl Tripp at ARUP Laboratories for tissue sectioning and IHC, and Drs. Stephan Ladisch, David Jones, and Linda Thompson for critically reading the manuscript, and Dr. Nikolaus S. Trede for his support at the project’s outset. An Institutional Development Award (IDeA) from the National Institute of General Medical Sciences (P20 GM103639) supports the OUHSC Stephenson Cancer Center’s Histology and Immunohistochemistry Core, which performed services for this project. Financial support: J.K.F. received support from Hyundai Hope On Wheels, the Oklahoma Center for the Advancement of Science and Technology (HRP-067), and an INBRE pilot project award from the National Institute of General Medical Sciences (P20 GM103447), and holds the E.L. & Thelma Gaylord Endowed Chair of the Children’s Hospital Foundation.

## Author contributions

C.B. and J.K.F. conceived and designed the research study. C.B., C.F., J.L.R., and J.K.F. analyzed the data. C.B., G.P., M.M., S.T.A., and J.B.G. performed experiments. T.S. and R.R.M. performed histologic analyses and imaging. G.t.K., J.L.R., and J.K.F. contributed essential reagents, tools and/or funding. S.B. and L.B. assisted with data analyses. C.B. and J.K.F wrote the manuscript. All authors revised and approved the final manuscript.

## Conflict of interest disclosure statement

Authors declare no conflict of interests.

## References

1. Jabbour E, O’Brien S, Konopleva M, and Kantarjian H. New insights into the pathophysiology and therapy of adult acute lymphoblastic leukemia. Cancer. 2015;121(15):2517–28.

2. Pui CH, Yang JJ, Hunger SP, Pieters R, Schrappe M, Biondi A, et al. Childhood Acute Lymphoblastic Leukemia: Progress Through Collaboration. J Clin Oncol. 2015;33(27):2938–48.

3. Woo JS, Alberti MO, and Tirado CA. Childhood B-acute lymphoblastic leukemia: a genetic update. Exp Hematol Oncol. 2014;3:16.

4. He S, Jing CB, and Look AT. Zebrafish models of leukemia. Methods Cell Biol. 2017;138:563–92.

5. Jing L, and Zon LI. Zebrafish as a model for normal and malignant hematopoiesis. Dis Model Mech. 2011;4(4):433–8.

6. Langenau DM, Traver D, Ferrando AA, Kutok JL, Aster JC, Kanki JP, et al. Myc-induced T cell leukemia in transgenic zebrafish. Science. 2003;299(5608):887–90.

7. Feng H, Langenau DM, Madge JA, Quinkertz A, Gutierrez A, Neuberg DS, et al. Heat-shock induction of T-cell lymphoma/leukaemia in conditional Cre/lox-regulated transgenic zebrafish. Br J Haematol. 2007;138(2):169–75.

8. Chen J, Jette C, Kanki JP, Aster JC, Look AT, and Griffin JD. NOTCH1-induced T-cell leukemia in transgenic zebrafish. Leukemia. 2007;21(3):462–71.

9. Frazer JK, Meeker ND, Rudner L, Bradley DF, Smith AC, Demarest B, et al. Heritable T-cell malignancy models established in a zebrafish phenotypic screen. Leukemia. 2009;23(10):1825–35.

10. Gutierrez A, Grebliunaite R, Feng H, Kozakewich E, Zhu S, Guo F, et al. Pten mediates Myc oncogene dependence in a conditional zebrafish model of T cell acute lymphoblastic leukemia. J Exp Med. 2011;208(8):1595–603.

11. Feng H, Stachura DL, White RM, Gutierrez A, Zhang L, Sanda T, et al. T-lymphoblastic lymphoma cells express high levels of BCL2, S1P1, and ICAM1, leading to a blockade of tumor cell intravasation. Cancer Cell. 2010;18(4):353–66.

12. Rudner LA, Brown KH, Dobrinski KP, Bradley DF, Garcia MI, Smith AC, et al. Shared acquired genomic changes in zebrafish and human T-ALL. Oncogene. 2011;30(41):4289– 96.

13. Blackburn JS, Liu S, Wilder JL, Dobrinski KP, Lobbardi R, Moore FE, et al. Clonal evolution enhances leukemia-propagating cell frequency in T cell acute lymphoblastic leukemia through Akt/mTORC1 pathway activation. Cancer Cell. 2014;25(3):366–78.

14. Blackburn JS, Liu S, Raiser DM, Martinez SA, Feng H, Meeker ND, et al. Notch signaling expands a pre-malignant pool of T-cell acute lymphoblastic leukemia clones without affecting leukemia-propagating cell frequency. Leukemia. 2012;26(9):2069–78.

15. Gutierrez A, Feng H, Stevenson K, Neuberg DS, Calzada O, Zhou Y, et al. Loss of function tp53 mutations do not accelerate the onset of myc-induced T-cell acute lymphoblastic leukaemia in the zebrafish. Br J Haematol. 2014;166(1):84–90.

16. Lobbardi R, Pinder J, Martinez-Pastor B, Theodorou M, Blackburn JS, Abraham BJ, et al. TOX Regulates Growth, DNA Repair, and Genomic Instability in T-cell Acute Lymphoblastic Leukemia. Cancer Discov. 2017;7(11):1336–53.

17. Ridges S, Heaton WL, Joshi D, Choi H, Eiring A, Batchelor L, et al. Zebrafish screen identifies novel compound with selective toxicity against leukemia. Blood. 2012;119(24):5621–31.

18. Gutierrez A, Pan L, Groen RW, Baleydier F, Kentsis A, Marineau J, et al. Phenothiazines induce PP2A-mediated apoptosis in T cell acute lymphoblastic leukemia. J Clin Invest. 2014;124(2):644–55.

19. Sabaawy HE, Azuma M, Embree LJ, Tsai HJ, Starost MF, and Hickstein DD. TEL-AML1 transgenic zebrafish model of precursor B cell acute lymphoblastic leukemia. Proc Natl Acad Sci U S A. 2006;103(41):15166–71.

20. Langenau DM, Ferrando AA, Traver D, Kutok JL, Hezel JP, Kanki JP, et al. In vivo tracking of T cell development, ablation, and engraftment in transgenic zebrafish. Proc Natl Acad Sci U S A. 2004;101(19):7369–74.

21. Anderson NM, Li D, Peng HL, Laroche FJ, Mansour MR, Gjini E, et al. The TCA cycle transferase DLST is important for MYC-mediated leukemogenesis. Leukemia. 2016;30(6):1365–74.

22. Cernan M, Szotkowski T, and Pikalova Z. Mixed-phenotype acute leukemia: state-of-the-art of the diagnosis, classification and treatment. Biomed Pap Med Fac Univ Palacky Olomouc Czech Repub. 2017;161(3):234–41.

23. Bauer TR Jr,., McDermid HE, Budarf ML, Van Keuren ML, and Blomberg BB. Physical location of the human immunoglobulin lambda-like genes, 14.1, 16.1, and 16.2. Immunogenetics. 1993;38(6):387–99.

24. Tang Q, Abdelfattah NS, Blackburn JS, Moore JC, Martinez SA, Moore FE, et al. Optimized cell transplantation using adult rag2 mutant zebrafish. Nat Methods. 2014;11(8):821–4.

25. Carmona SJ, Teichmann SA, Ferreira L, Macaulay IC, Stubbington MJ, Cvejic A, et al. Single-cell transcriptome analysis of fish immune cells provides insight into the evolution of vertebrate immune cell types. Genome Res. 2017;27(3):451–61.

26. Tang Q, Iyer S, Lobbardi R, Moore JC, Chen H, Lareau C, et al. Dissecting hematopoietic and renal cell heterogeneity in adult zebrafish at single-cell resolution using RNA sequencing. J Exp Med. 2017;214(10):2875–87.

27. Novershtern N, Subramanian A, Lawton LN, Mak RH, Haining WN, McConkey ME, et al. Densely interconnected transcriptional circuits control cell states in human hematopoiesis. Cell. 2011;144(2):296–309.

28. Haferlach T, Kohlmann A, Wieczorek L, Basso G, Kronnie GT, Bene MC, et al. Clinical utility of microarray-based gene expression profiling in the diagnosis and subclassification of leukemia: report from the International Microarray Innovations in Leukemia Study Group. J Clin Oncol. 2010;28(15):2529–37.

29. Moore FE, Garcia EG, Lobbardi R, Jain E, Tang Q, Moore JC, et al. Single-cell transcriptional analysis of normal, aberrant, and malignant hematopoiesis in zebrafish. J Exp Med. 2016;213(6):979–92.

30. Palmer C, Diehn M, Alizadeh AA, and Brown PO. Cell-type specific gene expression profiles of leukocytes in human peripheral blood. BMC Genomics. 2006;7:115.

31. Traver D, Paw BH, Poss KD, Penberthy WT, Lin S, and Zon LI. Transplantation and in vivo imaging of multilineage engraftment in zebrafish bloodless mutants. Nat Immunol. 2003;4(12):1238–46.

32. Zhang YA, Salinas I, Li J, Parra D, Bjork S, Xu Z, et al. IgT, a primitive immunoglobulin class specialized in mucosal immunity. Nat Immunol. 2010;11(9):827–35.

33. Malumbres R, Fresquet V, Roman-Gomez J, Bobadilla M, Robles EF, Altobelli GG, et al. LMO2 expression reflects the different stages of blast maturation and genetic features in B-cell acute lymphoblastic leukemia and predicts clinical outcome. Haematologica. 2011;96(7):980–6.

34. Liu X, Li YS, Shinton SA, Rhodes J, Tang L, Feng H, et al. Zebrafish B Cell Development without a Pre-B Cell Stage, Revealed by CD79 Fluorescence Reporter Transgenes.J Immunol. 2017;199(5):1706–15.

35. Dang CV. MYC on the path to cancer. Cell. 2012;149(1):22–35.

36. Meyer LH, Eckhoff SM, Queudeville M, Kraus JM, Giordan M, Stursberg J, et al. Early relapse in ALL is identified by time to leukemia in NOD/SCID mice and is characterized by a gene signature involving survival pathways. Cancer Cell. 2011;19(2):206–17.

37. Opgen-Rhein R, and Strimmer K. Accurate ranking of differentially expressed genes by a distribution-free shrinkage approach. Stat Appl Genet Mol Biol. 2007;6:Article9.

